# The structural basis for distinct binding avidity of Pertuzumab and Trastuzumab IgM towards HER2

**DOI:** 10.1101/709923

**Authors:** Firdaus Samsudin, Joshua Yi Yeo, Samuel Ken-En Gan, Peter J. Bond

## Abstract

Harnessing the therapeutic potential of immunoglobulin M (IgM) is of considerable interest in immunotherapy due to its complement-activating and cell-agglutinating abilities. Pertuzumab and Trastuzumab are monoclonal antibody drugs used in therapy for patients with human epidermal growth factor receptor 2 (HER2)-positive breast cancer but exhibit significantly different binding affinities as IgM when compared to the original IgG1 form. While the affinity of Pertuzumab IgM to the HER2 extracellular domain is about one order of magnitude higher than IgG1 in experiments, it was recently reported that Trastuzumab IgM and IgG have similar equilibrium dissociation constants to one another. We now perform an integrative multiscale simulation study in order to understand the structural basis for the differences in behavior between the two antibodies, based on complete antibody assemblies. We show that Pertuzumab IgM is able to utilize all of its V-regions to engage HER2 in a more stable mode than Trastuzumab IgM due to steric clashes between the large globular HER2 domains when bound to Trastuzumab. This is subsequently validated by confirming that Pertuzumab IgM inhibits proliferation in HER2 over-expressing live cells more effectively than its IgG1 counterpart. Given the widespread clinical use of Trastuzumab and Pertuzumab, elucidating the molecular details of antibody-antigen interaction may help guide the choice of epitopes for future design and selection of improved therapeutic antibody isotypes.

## INTRODUCTION

Immunoglobulin M (IgM) is the primary response antibody to combat foreign pathogens in adaptive immunity (Ehrenstein and Notley, 2010; Wang et al., 2016). As the first line of antibody defense, IgM tends to have lower antigen binding affinities. To compensate for this, secreted IgM forms multimeric structures (pentamers or hexamers), increasing the number of antigen binding sites for a higher overall avidity. This multimeric characteristic also confers other advantages to IgM. For example, the activation of the classical complement pathway requires the binding of multiple constant fragment (Fc) regions within close proximity, making multimeric IgM a very potent activator of the complement system (Cooper, 1985). The large size and multivalency of IgM molecules also enable the formation of bridges between distant epitopes, such as those on different viral particles, leading to superior aggregation properties when neutralizing viral infections (Baumgarth et al., 2000; Diamond et al., 2003; Seiler et al., 1991). Whilst most of the currently approved clinical monoclonal antibodies are of the IgG isotype, the high avidity of IgM and its effective complement activation and agglutination, makes IgM an attractive candidate for future immunotherapy (Kretschmer et al., 2017).

Multimeric IgM exists as either five (pentamer) or six (hexamer) subunits covalently linked to each other via disulfide bridges (Brewer et al., 1994; Davis and Shulman, 1989). Each IgM subunit is made of four polypeptide chains, namely two heavy chains containing five immunoglobulin (Ig) domains (Cμ1, Cμ2, Cμ3, Cμ4 and VH), and two light chains comprised of two Ig domains (CL and VL) (Figure S1). A short polypeptide called the joining (J)-chain may also be involved in IgM multimer formation, and the absence of the J-chain has been suggested to favor hexamer formation (Cattaneo and Neuberger, 1987; Niles et al., 2006). Due to the large size of the IgM pentamer and hexamer, high-resolution structural data for the entire complexes are absent. No crystal structure is currently available for the full-length monomeric IgM, let alone its pentameric or hexameric forms. Early studies based on negative-stain electron microscopy (EM) and small-angle X-ray scattering (SAXS) experiments suggested pentameric IgM to be a symmetric, star-shaped molecule with the antigen-binding fragment (Fab) regions pointing outwards (Davis et al., 1988; Feinstein and Munn, 1969; Perkins et al., 1991). Subsequently, cryo-atomic force microscopy (cryo-AFM) data showed the IgM pentamer to be non-planar, forming a mushroom-like shape with part of the Fc domains protruding out of the plane formed by the rest of the antibody (Czajkowsky and Shao, 2009). A model of IgM Fc was built based on SAXS analysis, integrating structures of each of the Cμ2, Cμ3, and Cμ4 domains solved using X-ray crystallography and NMR spectroscopy (Muller et al., 2013). Low-resolution cryo-electron tomography (cryo-ET) revealed that both the Fab and Fc domains of IgM are flexible and adopt multiple conformations (Akhouri et al., 2016). More recently, EM images indicated that in the presence of the J-chain, the IgM pentamer exhibits an asymmetric pentagonal shape with a large grove, acting as a carrier for apoptosis inhibitors in macrophages (Hiramoto et al., 2018).

Nevertheless, the structural details of interactions between multimeric IgM and antigens remain elusive, largely due to the experimental limitations associated with studying such large complexes. The molecular basis for how IgM achieves its strong avidity remains unclear. It is currently unknown whether all of the Fab domains in a multimeric IgM are able to bind antigens simultaneously, or whether the binding of an antigen on one Fab arm can affect the binding on another. The degree of avidity of IgM can also vary, especially when binding to different epitopes on the same antigen. For instance, the breast cancer therapeutic antibodies Pertuzumab and Trastuzumab (Baselga et al., 2012) showed remarkably different binding avidities to HER2 in their IgM form (Lua et al., 2018). Controlling for the different epitopes in HER2, there are significant discrepancies between the differences of Pertuzumab IgG1 and Trastuzumab IgG1 binding versus Pertuzumab IgM and Trastuzumab IgM. Compared to its monomeric IgG isotype counterpart, the equilibrium dissociation constants of Pertuzumab IgM to HER2 is around an order of magnitude higher. In contrast, Trastuzumab IgM had a similar equilibrium dissociation constant to HER2 for both IgG and IgM forms, suggesting a much weaker IgM avidity effect in the latter. A molecular-level understanding of how such distinct binding avidities arise for the same antigen is of importance for future design of therapeutic antibodies and epitope selection. Given that within the Pertuzumab and Trastuzumab models we have previously showed that the antibody-antigen interactions can be drastically affected by small changes in the antibody light chain (Su et al., 2017), antibody hinge (Su et al., 2018), V-region pairing (Ling et al., 2018), and VH families (Lua et al., 2019), it may be necessary to study the whole IgM molecule using a holistic approach, as previously proposed (Phua et al., 2019).

Thus, we now report the first integrative models of full-length Pertuzumab and Trastuzumab IgM multimers, based initially on available X-ray and NMR structures for each Ig domain. The models were validated against previously published EM and cryo-AFM data, while their structural stability and dynamics were assessed using a combination of extensive atomistic and coarse-grained (CG) molecular dynamics (MD) simulations. By modelling the binding of each antibody to the HER2 extracellular domain (ECD), we found that Pertuzumab IgM is able to form a geometrically optimal complex with its HER2 epitopes that enables multivalent interactions. On the other hand, Trastuzumab IgM binding to its HER2 epitope results in steric hindrances that inhibited simultaneous binding of multiple sites. Based on the Trastuzumab and Pertuzumab models, we show that IgM binding avidity depends primarily on the spatial location of the epitope on the antigen to enable efficient interaction with multiple sites at the same time. We subsequently validated this in a SKBR3 HER2 over-expressing breast cancer cell-line, where Pertuzumab IgM inhibited SKBR3 proliferation more effectively than Pertuzumab IgG1, whilst the reverse was observed for Trastuzumab. Collectively, our approach represents a novel strategy for providing guidance to epitope selection and antibody isotype selection in the future development of biologics.

## RESULTS

### Constructing Integrative Models of Pertuzumab and Trastuzumab IgM

We first built a homology model of a single unit (protomer) for each of Pertuzumab and Trastuzumab IgM (Figure 1A). The protomer comprises of four chains: two heavy and two light. The two heavy chains are covalently linked to each other via a disulfide bond between residues C331 in the Cμ2 domain. The heavy and light chains are linked via a disulfide bridge between residues C131 in the heavy chain and C214 in the C-terminus of the light chain. The structure of individual Ig domains for both antibodies have been resolved by X-ray crystallography and NMR spectroscopy (details in STAR Methods). IgM and IgE have a similar domain architecture within the Fc region with three Ig domains instead of two in the other antibody isotypes; we therefore used the structure of IgE Fc (Wan et al., 2002) to model the arrangement of Cμ2, Cμ3 and Cμ4 domains. A previous attempt at modelling the IgM pentamer also utilized the structure of IgE as a template due to its high sequence homology (Czajkowsky and Shao, 2009). As a result, our model displayed a sharp bend in the Fc part found in IgE.

**Figure 1.**
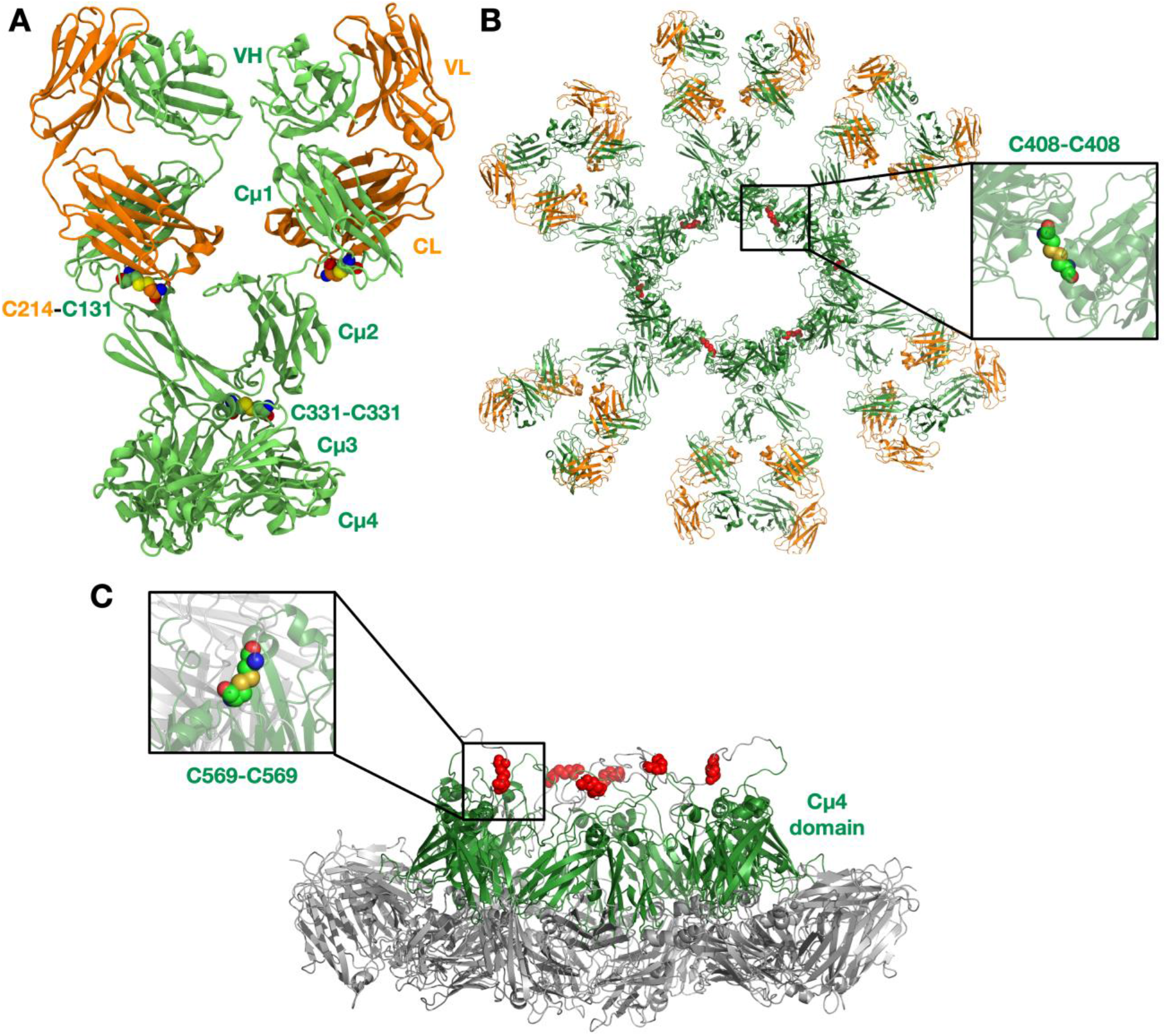
The Integrative model of Pertuzumab IgM. **(A)** Pertuzumab IgM protomer with inter-chain disulfide bridges highlighted. **(B)** Top view of Pertuzumab IgM hexamer. Disulfide bonds between the Cμ3 domain of adjacent IgM subunits are coloured in red and an example is highlighted in the inset. **(C)** Side view of the Pertuzumab IgM hexamer to show the protruding Cμ4 domains. Disulfide linkers between the C-terminal domains are illustrated in red and an example is highlighted in the inset. The Fab domains are not shown for clarity. IgM is shown in cartoon representation; the heavy and light chains are colored green and orange, respectively.

To test the structural integrity of the homology models, we performed 1 μs atomistic MD simulations, each system comprising ~300,000 atoms (Table 1). To assess the structural drift of each domain, we measured the root mean square displacement (RMSD) of the backbone atoms with respect to the initial structure. We found that for all Ig domains, the backbone RMSD reached a plateau after around 200 ns of 0.1-0.3 nm (Figure S2), suggesting that each domain was structurally stable. The radius of gyration (Rg) of each Ig domain also showed little fluctuation during the 1 μs simulations. Analysis of the secondary structure further corroborated this, showing a high degree of structural preservation for all Ig domains throughout (Figure S3). However, the backbone RMSD of the whole antibody was much higher, plateauing at ~1 nm in two simulations, but reaching as much as ~2.5 nm in the third. This indicates that the whole IgM protomer was structurally flexible such that each individual Ig domain could move relative to one another due to the unstructured linkers between them.

**Table 1:** List of Simulations.

| System contents | Resolution | Number of atoms / particles | Simulation Time (μs) |
| --- | --- | --- | --- |
| **Monomer** |  |  |  |
| Pertuzumab | Atomistic | 288,285 | 3 X 1 |
| Trastuzumab | Atomistic | 259,405 | 3 X 1 |
| Pertuzumab | Coarse-grained | 341,53 | 3 X 1 |
| Trastuzumab | Coarse-grained | 30,982 | 3 X 1 |
| **Hexamer** |  |  |  |
| Pertuzumab | Coarse-grained | 245,090 | 3 X 10 |
| Trastuzumab | Coarse-grained | 266,950 | 3 X 10 |
| Pertuzumab-HER2 | Coarse-grained | 559,507 | 3 X 10 |
| **Pentamer without J-chain** |  |  |  |
| Pertuzumab | Coarse-grained | 231,746 | 3 X 10 |
| Trastuzumab | Coarse-grained | 228,580 | 3 X 10 |
| **Pentamer with a J-chain** |  |  |  |
| Pertuzumab | Coarse-grained | 285,720 | 3 X 10 |
| Trastuzumab | Coarse-grained | 292,745 | 3 X 10 |

After validating the structural stability of the protomer domains, we built integrative models for IgM multimeric states. IgM exists either as pentamers or hexamers, and the former may be with or without the J-chain (a small ~130 amino acid residue protein involved in IgM assembly). IgM subunits within the oligomers are connected to one another via two disulfide linkages: one between residues C408 in the Cμ3 domains, and another between residues C569 in the C-terminal unstructured loops (Figure S1). In pentamers with a J-chain, the J-chain replaces the position of the sixth IgM subunit and connects to neighboring IgMs via disulfide bonds to residues C569. In this study, we built representative models for all three oligomeric states of IgM (Figure 1B and Figure S5). In agreement with cryo-AFM images (Czajkowsky and Shao, 2009), our resultant IgM oligomer models adopted a non-planar arrangement (Figure 1C). Due to the acute bend in the Fc region of the protomer, the Cμ4 domains and the C-terminal loop protrude out of the plane defined by the rest of the Fc domains. The J-chain in our pentameric model is also elevated with respect to the rest of the antibody as it is covalently linked to the C-terminal loop. Our model is also in agreement with a previous SAXS-derived model of the IgM Fc hexamer, where the Cμ4 domain formed a hexameric ring that formed the structural core of the whole antibody (Muller et al., 2013). Whilst the IgM hexamer and pentamer without a J-chain are symmetric, the pentamer with a J-chain formed an asymmetric pentagon as previously shown by single-particle negative-stain EM (Hiramoto et al., 2018). Due to spatial constraints, the Fab domains of one IgM subunit were necessarily positioned within close proximity to adjacent subunits, which could potentially have important implications for simultaneous antigen binding and, therefore, binding avidity.

### IgM Fabs are Flexible and Move Independently

IgM pentamers and hexamers are very large protein complexes (around 8,000 residues in the former and 10,000 residues in the latter), meaning that simulations at atomic resolution would be prohibitively expensive. As such, we next studied the dynamics of IgM pentamers and hexamers at coarse-grained (CG) resolution. An elastic network was applied to maintain the secondary structures within individual Ig domains of each IgM subunit. This elastic network was omitted between Ig domains as our atomistic simulations showed that individual domains may move freely relative to each other. We performed 1 μs CG simulations with a single IgM subunit and found that it showed similar flexibility to the atomistic model, with comparable distributions of relative domain-domain motions (Figure S4), showing our pseudo-atomic model to be a good approximation of the atomistic counterpart. The backbone RMSD and radius of gyration of each Ig domain throughout these simulations were also similar to the previously described atomistic simulations (Figure S2). To build CG models for IgM multimers, we also omitted the elastic network between adjacent IgM subunits; they are thus only linked covalently via disulfide bridges. Using these CG models, we then performed three independent repeats of 10 μs simulations per system.

We first looked at the structural dynamics of the IgM multimers. Comparing the per-residue root mean square fluctuation (RMSF) values as a measure of average flexibility along the chains, we found the Fc region to be rigidified due to the disulfide bridges linking each subunit (Figures 2 and S6). The Fab domains on the other hand were significantly more mobile with respect to their adjacent domains. This is consistent with negative-stain EM images of monoclonal mouse IgM pentamer, in which a strong density was observed for the Fc chains, whilst the peripheral region corresponding to the Fab domains could not be observed clearly due to their presumed flexible motion (Hiramoto et al., 2018). Principle component analysis (PCA) was used to assess collective motions for each complex. This revealed that the direction of motion of a given Fab domain is independent of its neighbors (Figure 3), suggesting allosteric regulation between IgM subunits to be unlikely. Due to their close proximity, adjacent Fab domains could also associate with one another via non-specific but long-timescale interactions, which reduced their mobility. These associations involved different Fab domains in different simulations, rationalizing the varying RMSF patterns of specific Fab domains observed across the three independent repeats (Figure S6).

**Figure 2.**
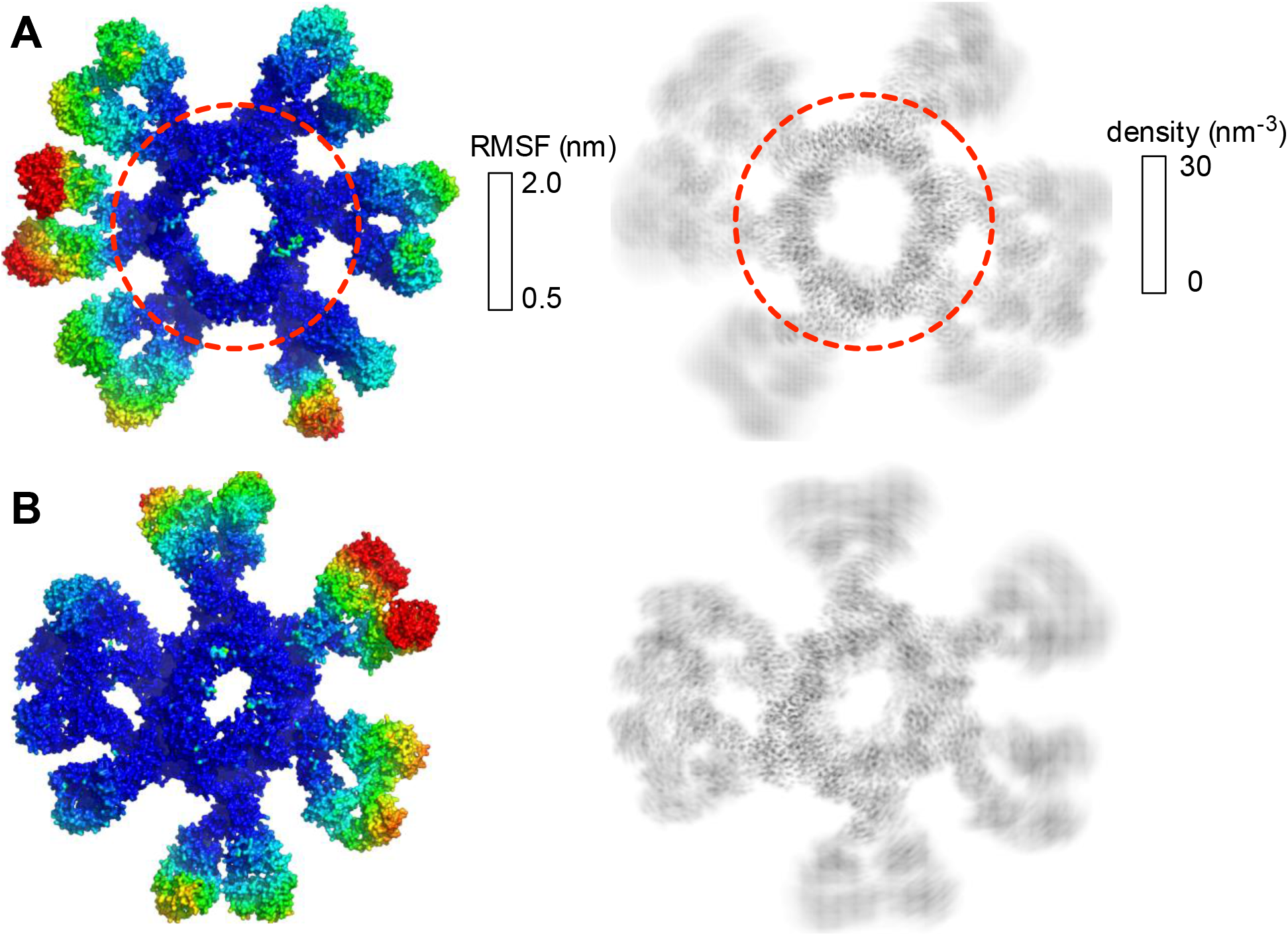
Flexibility of the Fab domain in hexameric IgM multimers. Root mean square fluctuation (RMSF) values were mapped onto the protein surface (left) and protein number density (right) from a 10 μs simulation for **(A)** Pertuzumab IgM hexamer and **(B)** Trastuzumab IgM hexamer. The dotted red line shows the approximate boundary between Fc and Fab domains.

**Figure 3.**
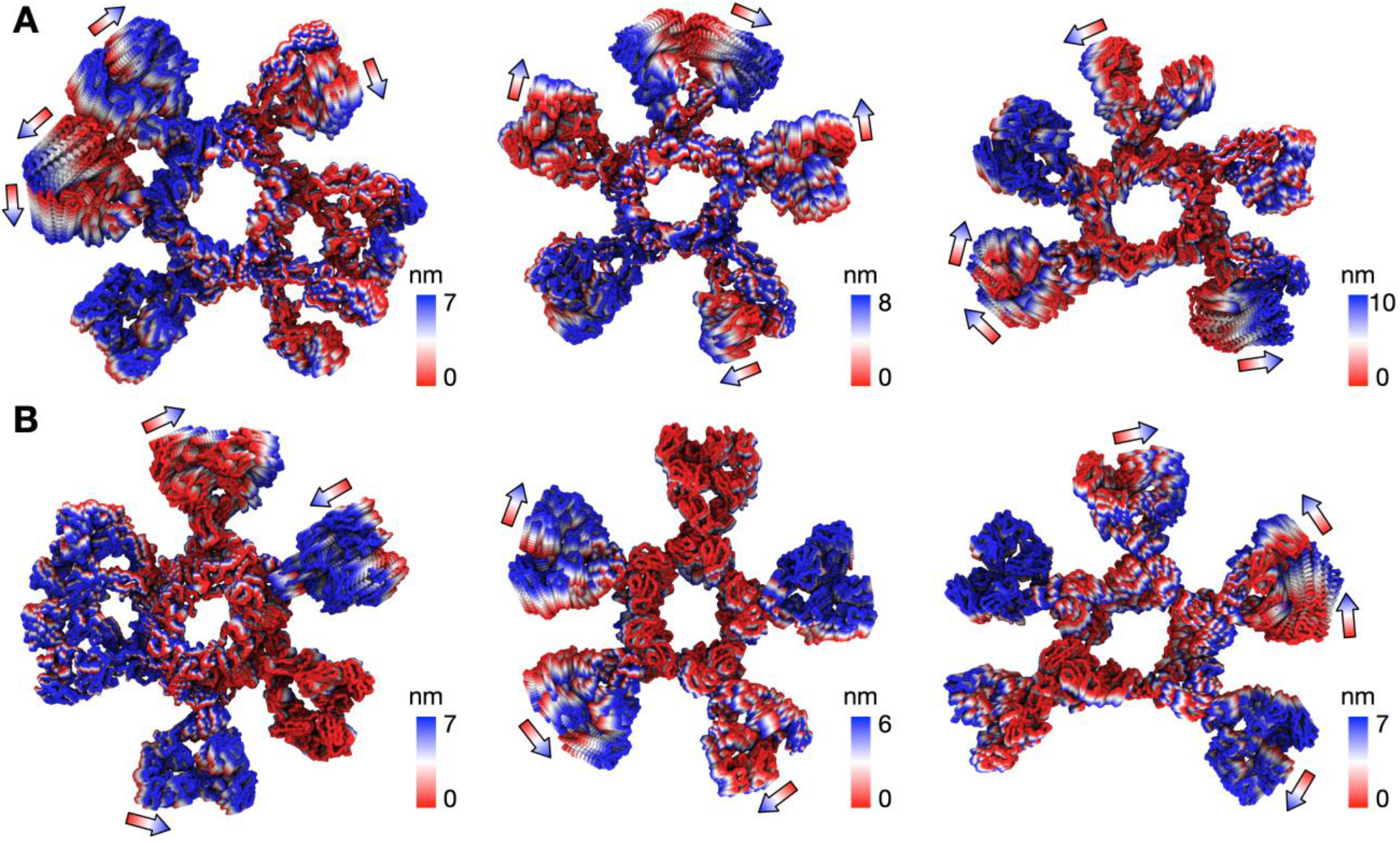
The direction of collective motion of the Fab domains during simulations based on principal component analysis (PCA). **(A)** The first principal motion of all backbone atoms as determined by PCA for: Pertuzumab IgM hexamer (left); Pertuzumab IgM pentamer without a J-chain (middle); and Pertuzumab IgM pentamer with a J-chain (right), each for one of the simulation replicas. The arrows illustrate how the direction of motion of the Fab domains is independent of adjacent neighbors. **(B)** The same analysis performed for simulations of Trastuzumab IgM multimers.

### Pertuzumab IgM and HER2 ECD form a Geometrically Optimal Complex

Avidity is an important factor for IgM given its ten or twelve antigen binding sites in its pentameric or hexameric forms, respectively. This contrasts with the only two or four binding sites found in other antibody isotypes. Although targeting the same HER2 antigen, Pertuzumab and Trastuzumab bind to different epitopes or sites on the HER2 ECD, and the effect of the different spatial arrangement of binding sites on the avidity of IgM is yet to be investigated. We therefore extended our models of the IgM hexamer and pentamer to integrate the bound HER2 ECD, based on previously determined crystal structures of monomeric Fab-HER2 complexes (Cho et al., 2003; Franklin et al., 2004).

Pertuzumab interacts with HER2 near the center of domain II found in the globular region of the ECD (Figure S7). This blocks the dimerisation hairpin from fitting into its binding site on adjacent receptors. To build a model of HER2 bound Pertuzumab IgM, we performed structural alignment of each of the Fab domains in the initial integrative model of the Pertuzumab IgM hexamer with the crystal structure (Figure 4). This model revealed the exposed region, primarily on domain IV, to be small and free of steric clashes. This makes it possible to fit twelve HER2 ECDs to the IgM hexamer, one for each Fab, maximizing the multiple binding sites on IgM, expectedly increasing avidity.

**Figure 4.**
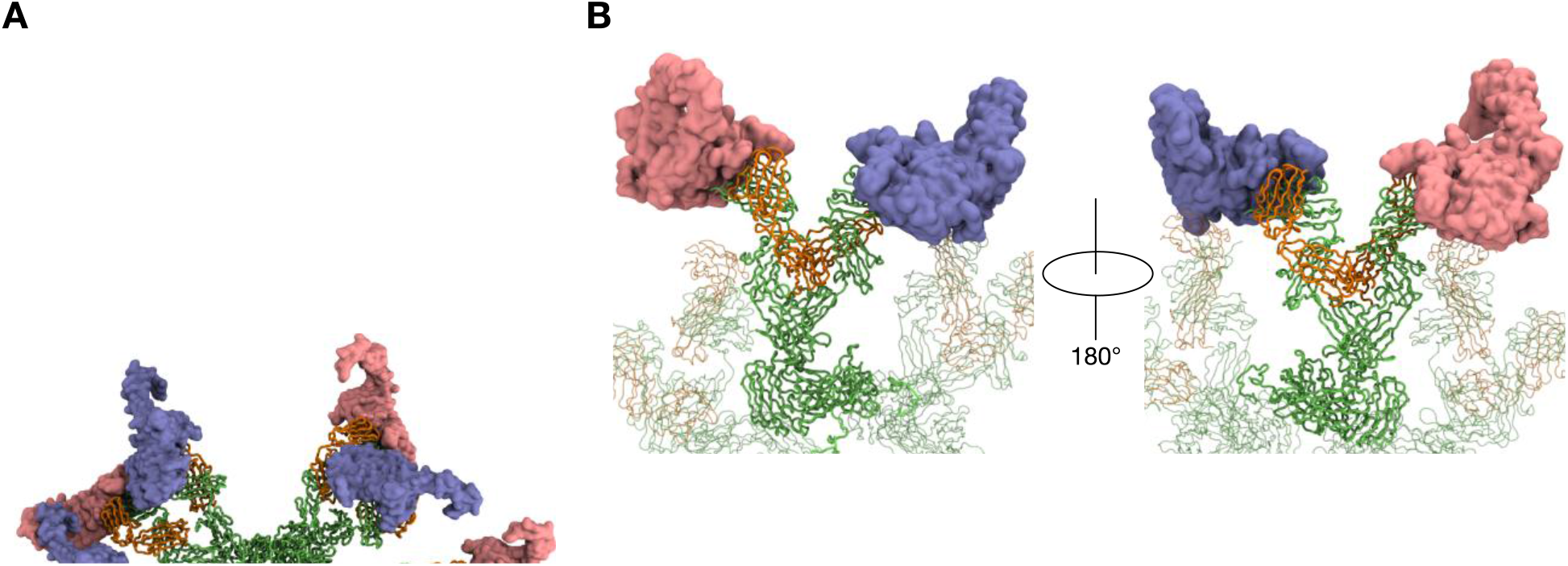
The binding of HER2 ECD to Pertuzumab IgM hexamer. **(A)** Twelve HER2 ECDs (pink and purple) were aligned onto Pertuzumab IgM hexamer initial model, one for each Fab domain, based on the previously determined binding site (Franklin et al., 2004). **(B)** Enlarged images of one of the IgM subunits with HER2 ECDs bound. The rest of the IgM is shown in transparent representation and the other HER2 ECDs are not shown for clarity. IgM is shown in ribbon representation and coloured green and orange for heavy and light chains, respectively, and HER2 ECDs are shown in surface representation.

To test the binding stability, we performed three independent 10 μs simulations of the Pertuzumab IgM hexamer complexed to multiple HER2 ECDs. In all three simulations, all twelve HER2 ECDs remained bound to the antibody; the RMSD of the Fab-HER2 complexes reached a plateau at around 0.5 nm, suggesting stable binding (Figure 5). While the globular region of HER2 ECD remained relatively rigid due to the binding to IgM, the solvent exposed domain IV was more flexible. To understand the effect of HER2 binding on the conformational flexibility of Pertuzumab IgM, we measured its per-residue RMSF. Similar to the apo simulations, the Fab regions exhibited more motion compared to the Fc part. Interestingly, the RMSD of the Fc domain from all three HER2-bound simulations was comparable to that of the apo simulations, suggesting that HER2 binding did not affect the conformation of IgM Fc.

**Figure 5.**
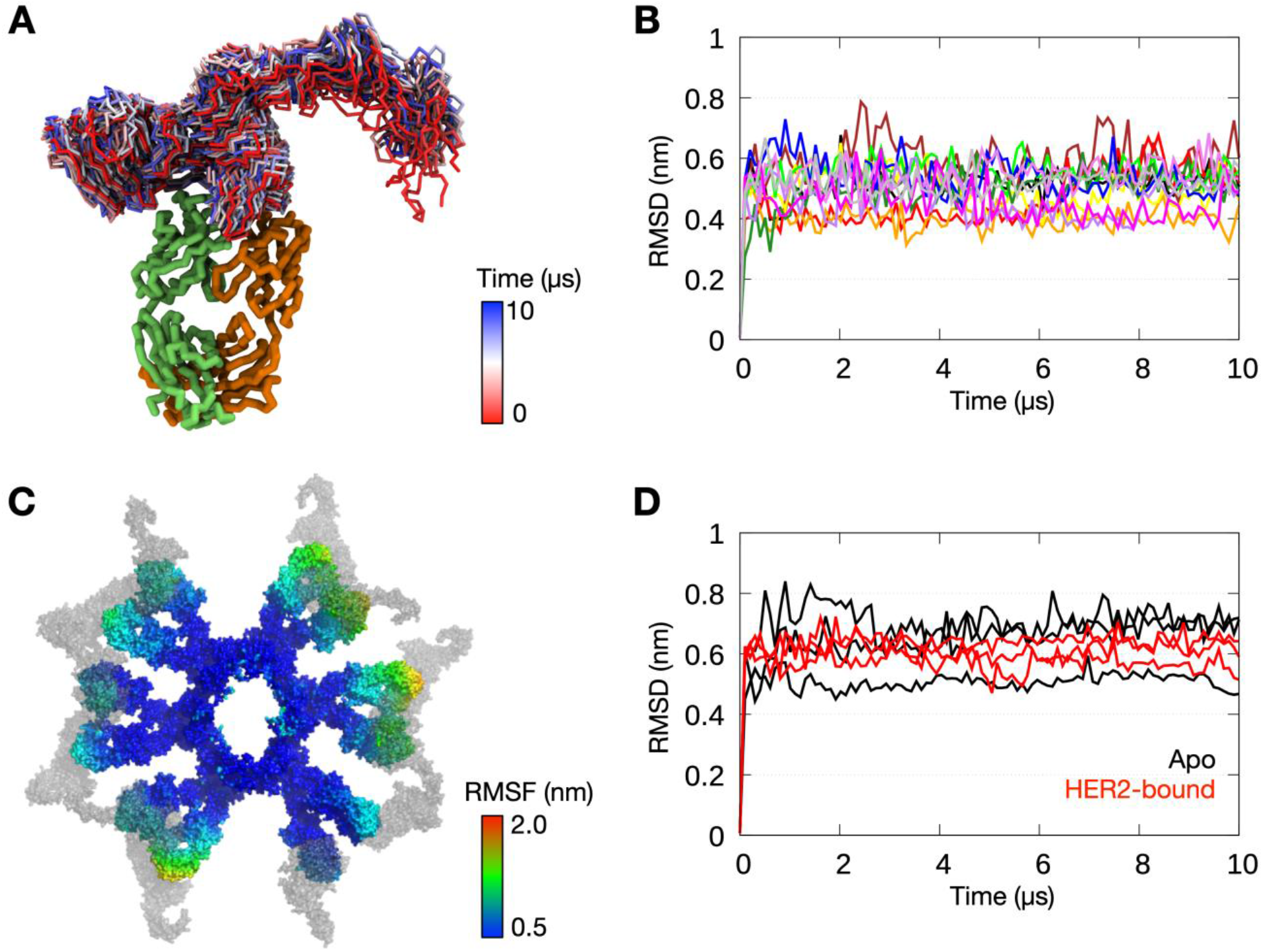
Simulations of Pertuzumab IgM hexamer complexed with HER2 ECDs. **(A)** Conformations sampled by one of the HER2 ECDs during one of the 10 μs simulations. The bound Fab domain is shown in green (heavy chain) and orange (light chain), and equally spaced conformations of the ECD are shown overlaid, colored according to timeframe. Only the backbone atoms are shown for clarity. **(B)** Backbone RMSDs of each of the twelve Fab-HER2 complexes during the simulation showed stable binding. **(C)** Per-residue RMSF of the Pertuzumab IgM hexamer during one of the 10 μs simulations. HER2 molecules are shown in grey. **(D)** Comparison of backbone RMSDs of the Fc domain of IgM hexamer from apo versus HER2-bound simulations. Data is shown for three independent repeats of each system.

### Steric Clashes Hinder Multiple Binding of HER2 to Trastuzumab IgM

In contrast with Pertuzumab, Trastuzumab binds HER2 on the C-terminal portion of domain IV, close to its transmembrane domain (Cho et al., 2003). This binding poses a steric barrier for optimal interaction between the transmembrane regions of neighboring HER2 receptors, which is required for signaling. As performed above, we placed the HER2 ECD onto the initial Trastuzumab IgM hexamer model using structural alignment based on crystallographic data (Figure 6). We found that the binding of one HER2 ECD to a Fab domain resulted in significant steric clashes to the adjacent Fab domain on the same IgM subunit. As Trastuzumab interacts with the small juxtamembrane region of HER2, the large globular region composed of domains I, II, and III is exposed. Therefore, for HER2 binding to occur, adjacent Fab domains must have sufficient space between them to accommodate this globular region of HER2 ECD.

**Figure 6.**
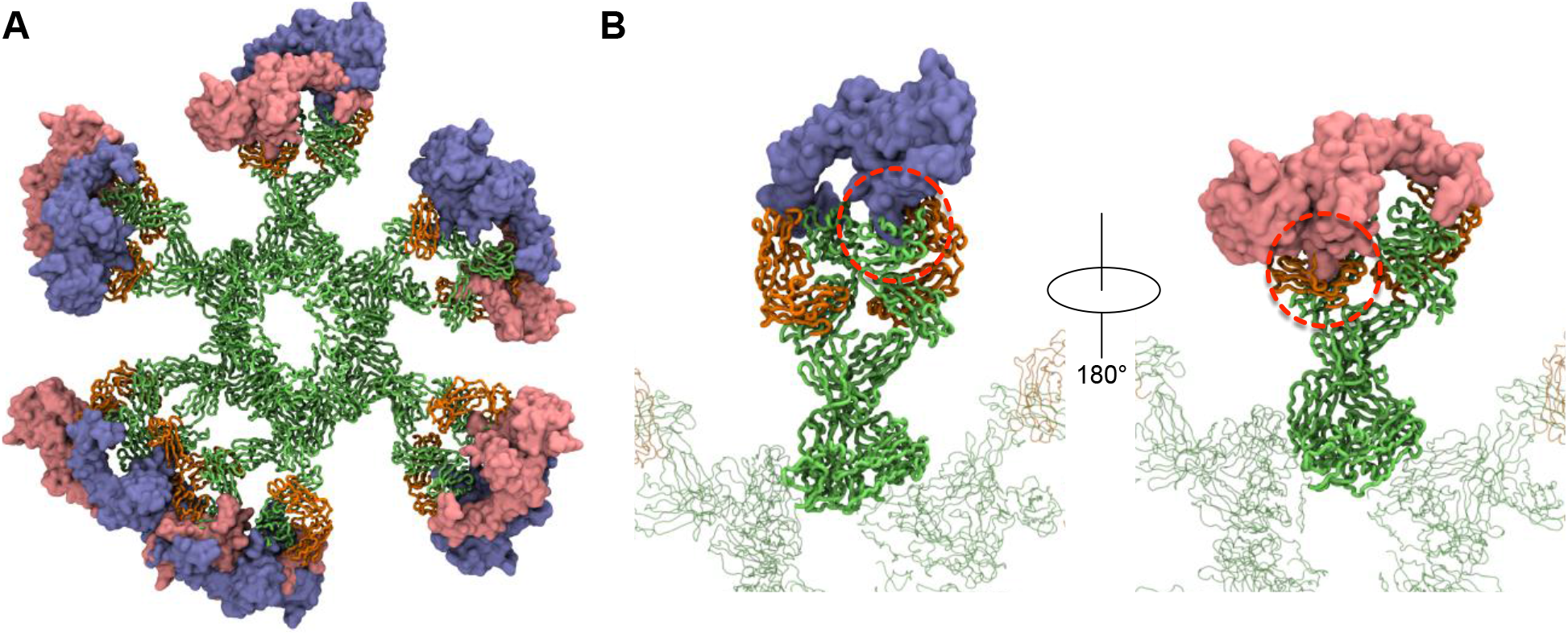
The binding of HER2 ECD to Trastuzumab IgM hexamer. **(A)** Twelve HER2 ECDs (pink and purple) aligned onto the initial model of Trastuzumab IgM hexamer based on the previously elucidated monomeric binding site (Cho et al., 2003). **(B)** Enlarged images of one IgM subunit with HER2 bound. Dotted red line highlights the steric clashes between the globular domain of HER2 and the adjacent Fab domain of IgM. Only one HER2 ECD is shown in each image for clarity. IgM is shown in ribbon representation and coloured green and orange for heavy and light chains, respectively, and HER2 ECDs are shown in surface representation.

As the IgM Fab is flexible, it is possible that adjacent Fab domains may move apart from one another to allow HER2 binding. To approximate the likelihood of HER2 binding from our simulations, we first estimated the distance a Fab domain has to move with respect to its neighbor to accommodate HER2. This was performed by measuring the width of the exposed globular domain of HER2 from the crystal structure. We found that a clearance of around 1.2 nm and 1.5 nm on either side of HER2 binding site would allow for binding without a steric clash (Figure 7A). We then measured distances along the same axis between adjacent Fab domains from our apo simulations to determine how frequently this configuration was achieved. Our apo simulations were 10 μs each with snapshots recorded every 1 ns, giving a total of 10,000 frames. As there are twelve or ten Fabs in IgM hexamer or pentamer, respectively, there were 300,000-360,000 Fab configurations per triplicate trajectory for each system. The resultant distribution of distances is shown in Figure 7B. For the Trastuzumab IgM hexamer, the distance requirement to allow HER2 binding was only found ~500 times, giving rise to a probability of around 1.4 × 10^−3^. This probability was slightly higher for simulations of IgM pentamers with and without a J chain, with a probability of around 1.7 × 10^−2^ and 3.7 × 10^−2^, respectively. Overall, these results further corroborate the low probability of simultaneous binding of HER2 ECD to Trastuzumab IgM, helping to rationalize the weaker avidity effect compared to Pertuzumab.

**Figure 7.**
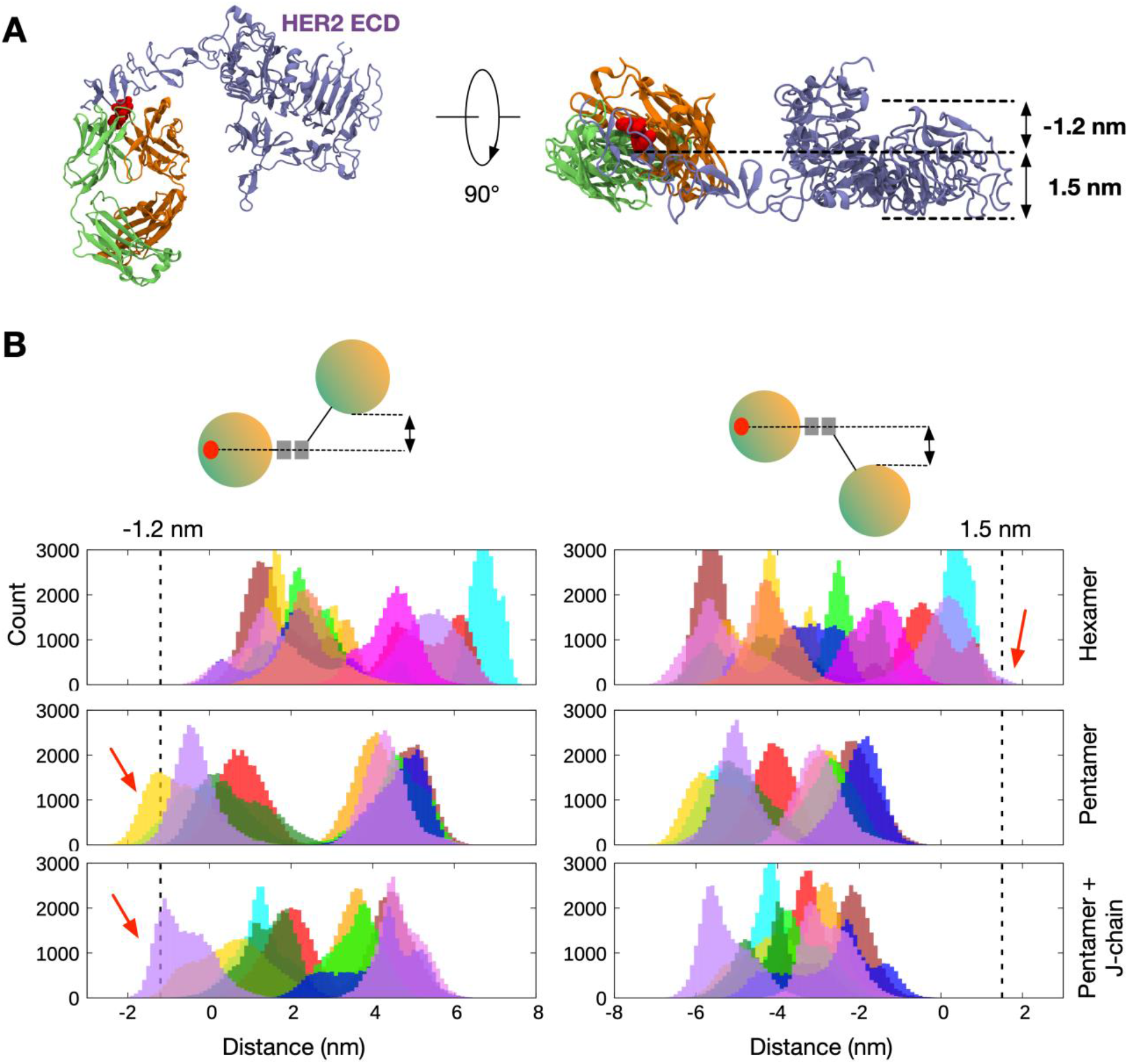
The distribution of distances between adjacent Fab domains from CG simulations of Trastuzumab IgM. **(A)** Side (left) and top (right) views of the crystal structure of Trastuzumab Fab-HER2 ECD complex. Light and heavy chains are orange and green, respectively, while the binding site residues are highlighted in red. The space required to accommodate HER2 ECD is shown on the right. **(B)** Distance distribution between the center of mass of the binding site residues on a Fab domain and the bottom-most (left) or the top-most (right) point on the adjacent Fab domain. Different colors depict the ten/twelve different Fab domains in each IgM. The data was extracted from three independent apo simulations of Trastuzumab IgM hexamer, pentamer, and pentamer with a J-chain. Dotted lines indicate the distance cut-off required for HER2 binding and red arrows show the parts of the distance distributions that meet this requirement.

### Pertuzumab IgM Inhibits Cell Proliferation more Effectively than IgG1

Finally, we sought to validate our models by testing the inhibitory effectiveness of the Pertuzumab and Trastuzumab IgM and IgG1 in live cells. Thus, using HER2 over-expressing breast cancer cell line SKBR3, cells were grown in the presence of Pertuzumab or Trastuzumab in their IgM or IgG1 forms, or a water control, for 7 days, before counting the viable cells. Even though Pertuzumab and Trastuzumab IgM and IgG1 inhibited cell growth compared to the control, Pertuzumab IgM inhibited cell proliferation significantly more effectively than Pertuzumab IgG1, whereas the reverse was found for Trastuzumab IgM (Figure 8), thus supporting the hypothesized inhibitory mechanism based on analysis of our models.

**Figure 8:**
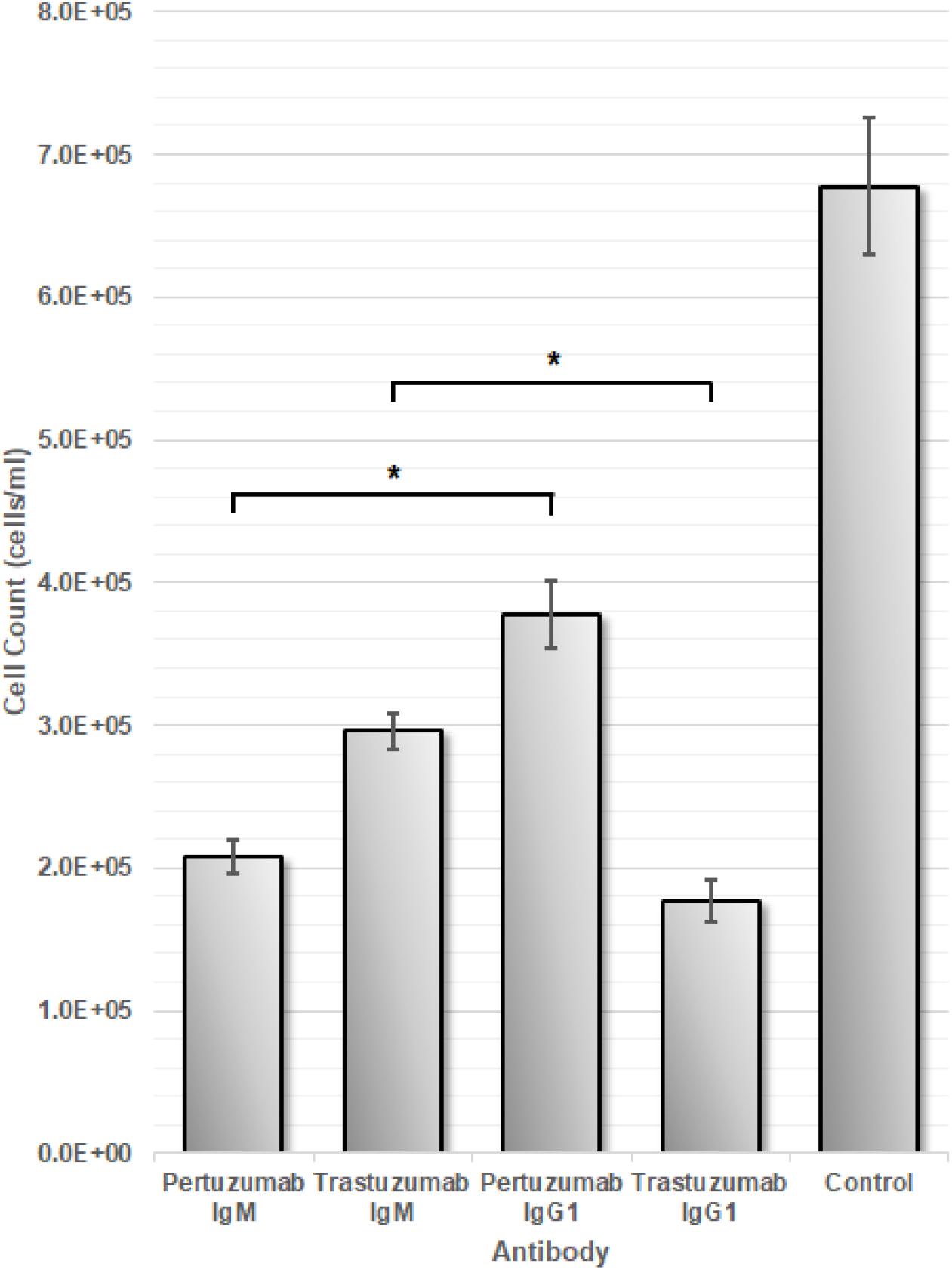
SKBR3 Cell Count Assay. HER2-positive SKBR3 cells were incubated with one of four antibodies (Pertuzumab IgM, Trastuzumab IgM, Pertuzumab IgG1, Trastuzumab IgG1) or a PBS (control). Cell count (cells/ml) of the culture was performed 7 days after incubation with the antibody. The error bars are the differences from three independent batches of the cell assay. * represents statistical significance (p < 0.05).

## DISCUSSION

We set out to report full-length models of multimeric IgM antibodies that matched existing experimental data, followed by multiscale simulations to study the detailed molecular mechanism of IgM antigen binding avidity. Our models agreed with previously published structural data and showed good overall structural integrity during simulations. We found the Fab domains in multimeric IgM to be flexible whilst the Fc domains were rigid due to the interconnecting disulfide bonds between different subunits. Each Fab arm moved independently of its neighbors suggesting that long-distance allostery between different IgM subunits is unlikely. Models of HER2 binding to IgM hexamers revealed that Pertuzumab could bind to twelve copies of HER2 ECD simultaneously, explaining the impressive avidity effect compared to its monomeric isotypes (Lua et al., 2018). Trastuzumab IgM hexamers, however, could not bind to multiple copies of HER2 due to the steric clashes caused by the large globular domain of the latter. The hypothesized inhibitory mechanism for each antibody was then validated by testing their effectiveness in live cells.

There are two possible caveats in our study. Firstly, the use of an elastic network model in our CG simulations limits possible large conformational changes in the antibodies. While the Fab arms displayed some degree of flexibility in our simulations, we did not observe any dramatic bending in the Fc part. This behaviour contrasts with cryo-ET structures of IgM, which indicated that the core Fc region could distort resulting in a bell-shaped conformation (Akhouri et al., 2016), but is in good agreement with negative-stain EM data (Hiramoto et al., 2018). Secondly, we have not explored the possible effects of glycosylation in our model due to the complexity of generating parameters for the CG simulations of polysaccharides. Most of IgM glycosylation sites, however, are found within the Fc region (Arnold et al., 2005), and are thereby unlikely to affect HER2 binding at the Fab domains. Earlier simulation studies showed that N-glycosylation did not trigger large changes in the overall protein structure but rather reduced protein dynamics (Lee et al., 2015). This may aggravate simultaneous binding of HER2 in the case of Trastuzumab IgM binding.

Allosteric modulation has been an area of interest in molecular immunology (Janda et al., 2016; Lua et al., 2019; Su et al., 2017; Yang et al., 2017). Previous experimental and computational studies of IgG and IgA have suggested that the Fc and Fab regions of an antibody might communicate with one another (Ling et al., 2018; Phua et al., 2019); for example, point mutations on the Fc region may reduce binding affinity, whilst antigen binding to the Fab region changes the conformation of the Fc domain to promote binding to its receptor (Lua et al., 2019; Oda et al., 2003; Schlessinger et al., 1975; Su et al., 2018). Allostery between the different subunits in a multimeric IgM, however, remain elusive. Our CG simulations indicated that the individual Fab domains are free to move independently of one another and do not affect the motion of their neighbors. Furthermore, we did not observe any difference in the flexibility of the Fc region upon binding with HER2 compared to the apo systems within the timescale of our simulations, suggesting that allosteric communication between one Fab domain and another via the Fc region is unlikely to occur. As previously discussed, the use of an elastic network model also limits any large conformational changes that may be needed for allostery. Nonetheless, allosteric modulation did not need to be invoked in order to rationalize the experimentally validated HER2 inhibitory activity of each antibody.

Our study highlights the importance of studying the antibody-antigen binding interface in the context of a biologically realistic assembly, specifically in multivalent isotypes like IgM with Fab domains in close proximity to one another. To date, therapeutic antibodies are mostly of the bivalent isotypes, in particular IgG; however, studies on using IgM for clinical purposes are underway especially for cancer treatment (Brandlein et al., 2007; Liedtke et al., 2012; Rasche et al., 2013, 2015). This is unsurprising given that IgM is immunologically superior than other isotypes, vis-à-vis antigen agglutination and complement activation. The dramatic increase in overall antigen binding strength due to avidity effects is another compelling reason to use IgM. Previous experimental measurement of binding affinities involving IgM and other monomeric isotypes, however, showed that the overall increase in binding avidity in the former is not always significant compared to the latter (Lua et al., 2018). This suggests that the addition of antigen binding sites in multimeric IgM would not guarantee an improved overall binding. Indeed, our modelling and simulations show that Trastuzumab IgM is not capable of utilizing all of its Fab domains for HER2 binding due to steric clashes, which explains its poorer avidity compared to Pertuzumab IgM. Key to this steric barrier is the Pertuzumab-HER2 binding interface, located at the juxtamembrane region of the latter (Cho et al., 2003). We postulate that in the presence of the cell membrane and other membrane proteins *in vivo*, Trastuzumab IgM may be severely hindered from accessing HER2 juxtamembrane regions due to its large size, rendering it a less effective choice compared to the smaller IgG as shown by the SKBR3 assay where the Trastuzumab IgM was significantly less effective at inhibiting SKBR3 proliferation than Trastuzumab IgG1. On the other hand, Pertuzumab IgM is able to bind HER2 in all of its antigen binding sites without any steric clashes due to the binding interface being positioned at the large globular domain of HER2 (Franklin et al., 2004). This capability is especially beneficial to elicit multiple simultaneous interactions in cancer cells overexpressing HER2 receptors. As such, the Pertuzumab IgM isotype may be more effective than its IgG counterpart for HER2+ cancer therapy as also shown in the SKBR3 assay. Our results shed light on the crucial role of understanding antibody-antigen interactions at the molecular level for more effective antibody isotype selection, as well as epitope selection for biologics development.

## ACKNOWLEDGEMENTS

This work was supported by A*STAR BII APD Lab core funds. Simulations were performed on the petascale computer cluster ASPIRE-1 at the National Supercomputing Center of Singapore.

## AUTHOR CONTRIBUTIONS

Conceptualization, S.K-E.G, and P.J.B.; Computational investigation, F.S.; Wet lab investigation, J.Y.Y.; Writing – Original Draft, F.S., S.K-E.G and P.J.B.; Supervision, S.K-E.G and P.J.B.

## STAR METHODS

### CONTACT FOR RESOURCE SHARING

Further information and requests for resources should be directed to and will be fulfilled by the Lead Contact, Peter J. Bond. Trastuzumab and Pertuzumab IgM and IgG1 proteins can be requested from Samuel Gan

## METHOD DETAILS

### Modelling IgM Hexamers and Pentamers

The homology models of Pertuzumab and Trastuzumab IgM protomers were constructed using Modeller version 9.21 (Sali and Blundell, 1994). The crystal structures of the Fab domains of Pertuzumab and Trastuzumab in complex with human HER2 ECD (PDB: 1S78 (Franklin et al., 2004) and 1N8Z (Cho et al., 2003), respectively) were used as the template for the Fab region, whilst the Fc region was modelled based on the atomic structures of its three domains: Cμ2 (PDB: 4JVU), Cμ3 (PDB: 4BA8) and Cμ4 (PDB: 4JVW) previously determined using X-ray crystallography and NMR spectroscopy (Muller et al., 2013). For each antibody, 10 models were built and the model with the lowest discreet optimized protein energy (DOPE) score (Shen et al., 2006) was chosen. Stereochemical assessment using Ramachandran analysis (Ramachandran et al., 1963) of the non-loop regions showed 40 and 31 outlier residues for Pertuzumab and Trastuzumab IgM models, respectively, representing around 2% of the whole protein, confirming that the models are structurally reasonable. Further structural assessment was based on analysis of stability during all-atom MD simulations. To build models of the hexameric and pentameric IgM, the protomers were duplicated and aligned within PyMOL (DeLano, 2002) and two disulfide bridges were constructed between adjacent protomers (Figure S1). For the IgM pentamer with a J-chain, the 137-residue J-chain (UniProt: P01591) was modelled as an unstructured loop as there is currently no crystallographic data available. Disulfide bonds within the J-chain and between the J-chain and neighboring IgM protomers were constructed as reported by Frutiger et al. (Frutiger et al., 1992); these disulfide bridges maintained the rigidity of the J-chain as shown by the similar RMSF values compared to the Fc domains in the pentamer. To model the binding of HER2 ECD to Pertuzumab and Trastuzumab IgM hexamers, we aligned the respective HER2-Fab complexes to each of the Fab domains onto the initial hexameric models. Missing residues in HER2 ECD were built as loops using Modeller.

### Simulation Systems and Protocols

Atomistic MD simulations were performed to test the structural stability of Pertuzumab and Trastuzumab IgM protomers. The antibodies were parameterized using CHARMM36 force field (Huang and MacKerell, 2013) and solvated in TIP3P water molecules with 0.15 M NaCl added to neutralized the system. Short 100 ps equilibration simulations were performed during which the heavy atoms of the antibodies were positionally restrained using a force constant of 1000 kJ mol^−1^ nm^−2^. The temperature was maintained at 310 K using the Nose-Hoover thermostat with a time constant of 1.0 ps (Hoover, 1985; Nosé, 1984), while the pressure was kept at 1 atm by an isotropic coupling to the Parrinello-Rahman barostat with a time constant of 5.0 ps (Parrinello, 1981). The smooth particle mesh Ewald method with a real-space cut-off of 1.2 nm was utilized to calculate the electrostatic interactions (Essmann et al., 1995). The Van der Waals interactions were truncated at 1.2 nm with a force switch smoothing function applied from 1.0 to 1.2 nm. All covalent bonds were constrained using the LINCS algorithm (Hess et al., 1997), which allowed for an integration time step of 2 fs. After the equilibration simulations, the positional restraints were removed and three independent 1 μs production simulations were conducted, each starting with a different distribution of atomic velocities.

To understand the dynamics of Pertuzumab and Trastuzumab IgM hexamers and pentamers with and without HER2 ECD, we performed coarse-grained (CG) MD simulations. Antibodies and HER2 ECD were converted to CG representation using the MARTINI 2.2 force field with elastic network ElNeDyn model applied to maintain the secondary structures (Monticelli et al., 2008; Periole et al., 2009). The systems were solvated using standard MARTINI water molecules and neutralized using 0.15 M NaCl. Equilibration simulations were performed for 10 ns whereby positional restrains with a force constant of 1000 kJ mol^−1^ nm^−2^ was applied to the backbone atoms of the proteins. The temperature was maintained at 310 K using a velocity-rescaling thermostat with a time constant of 1 ps (Bussi et al., 2007), while the pressure was kept at 1 atm using an isotropic coupling to a Berendsen barostat with a time constant of 5 ps (Berendsen et al., 1984). The electrostatics were computed using the reaction field method, whilst the Lennard-Jones potential was cut off using a potential shift Verlet scheme; the short-range cut-off for both was set to 1.1 nm. Following the equilibration simulations, three independent 10 μs simulations were conducted with different starting velocities. For these production runs, the Parrinello-Rahman barostat was utilized with a coupling constant of 12.0 ps (Parrinello, 1981). The list of simulations performed for this study is available in Table 1. All simulations were performed using GROMACS package version 2018 (Abraham et al., 2015a) and visualized in PyMOL (DeLano, 2002) and VMD (Humphrey and Dalke, 1996).

### Analysis

All analysis was performed using GROMACS package version 2018 (Abraham et al., 2015b) or custom Python scripts utilizing the MDAnalysis toolkit (Michaud-agrawal et al., 2011).

### Inhibition of Proliferation of HER2-Positive SKBR3

HER2-positive *SKBR3* cell line was maintained in T-25 flasks. Cells were cultured in DMEM (Gibco) and seeded into Corning® Costar® 6-well cell culture plates (Costar) at 2 × 10^5^/ mL. Respective antibodies previously produced (Pertuzumab IgM, Trastuzumab IgM, Pertuzumab IgG1, Trastuzumab IgG1, see (Lua et al., 2018)) were added at 1 μg/ml. After 7 days, viable cells were counted using a haemocytometer with Trypan Blue stain 0.4% (Invitrogen). Paired sample t-test was utilized to perform statistical analysis.

## QUANTIFICATION AND STATISTICAL ANALYSIS

See **METHOD DETAILS** above.

**KEY RESOURCES TABLE.**
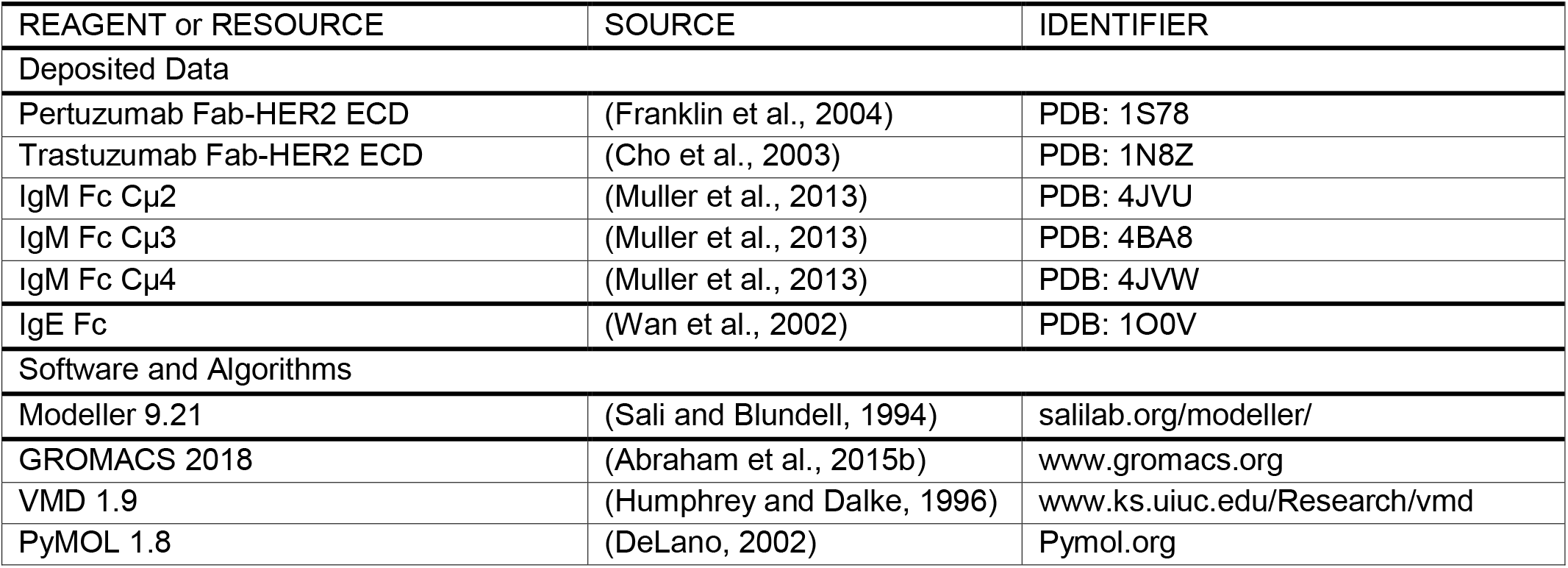

