## Supplementary Materials for "The structural basis for distinct binding avidity of Pertuzumab and Trastuzumab IgM towards HER2"

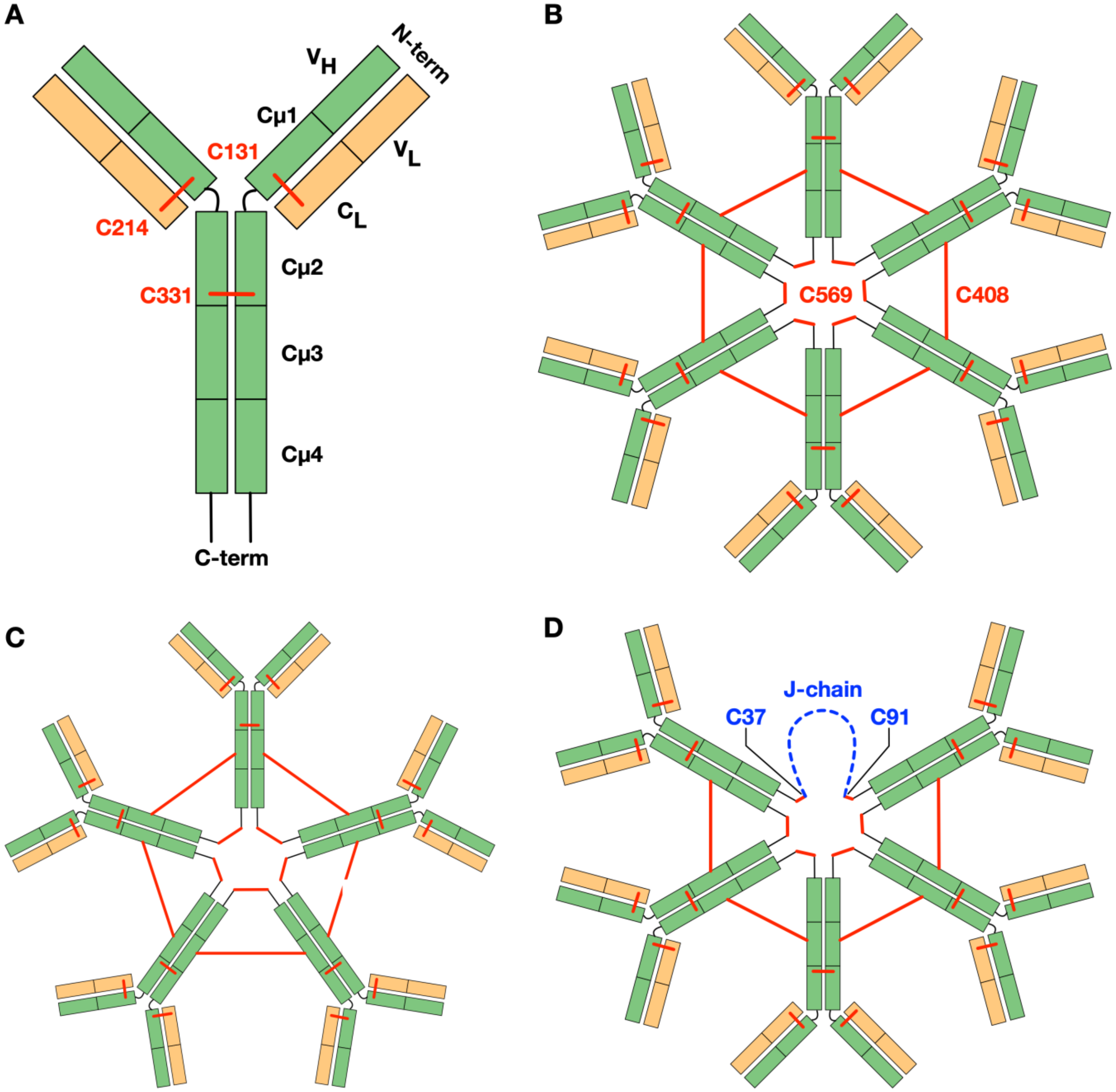


**Figure S1 expositing Figure 1: Schematics of IgM and the positions of interchain disulfide bridges. (A)** The IgM protomer consists of two heavy chains (green) and two light chains (orange). The former is made of five immunoglobulin (Ig) domains (heavy chain constant domains Cμ1-4 and heavy chain variable domain VH), whilst the latter is comprised of two domains: light chain constant (CL) and light chain variable (VL). The two heavy chains are covalently linked by a disulfide bond between the Cμ2 domains, and the light chains are connected to adjacent heavy chains via a disulfide bond in their C-termini. **(B)** IgM hexamers consists of six units of IgM, each linked to another via two disulfide bonds: one on the Cμ3 domains and the other on the C-terminus of the heavy chain. **(C)** IgM pentamers consists of five constant regions of IgM, which are connected to each other via the same disulfide bridges as the hexamer. **(D)** Pentameric IgM with a J-chain is an asymmetric pentagon, whereby the J-chain replaces the sixth IgM constant region and connects to the neighboring IgM via disulfide bonds on their C-terminus.

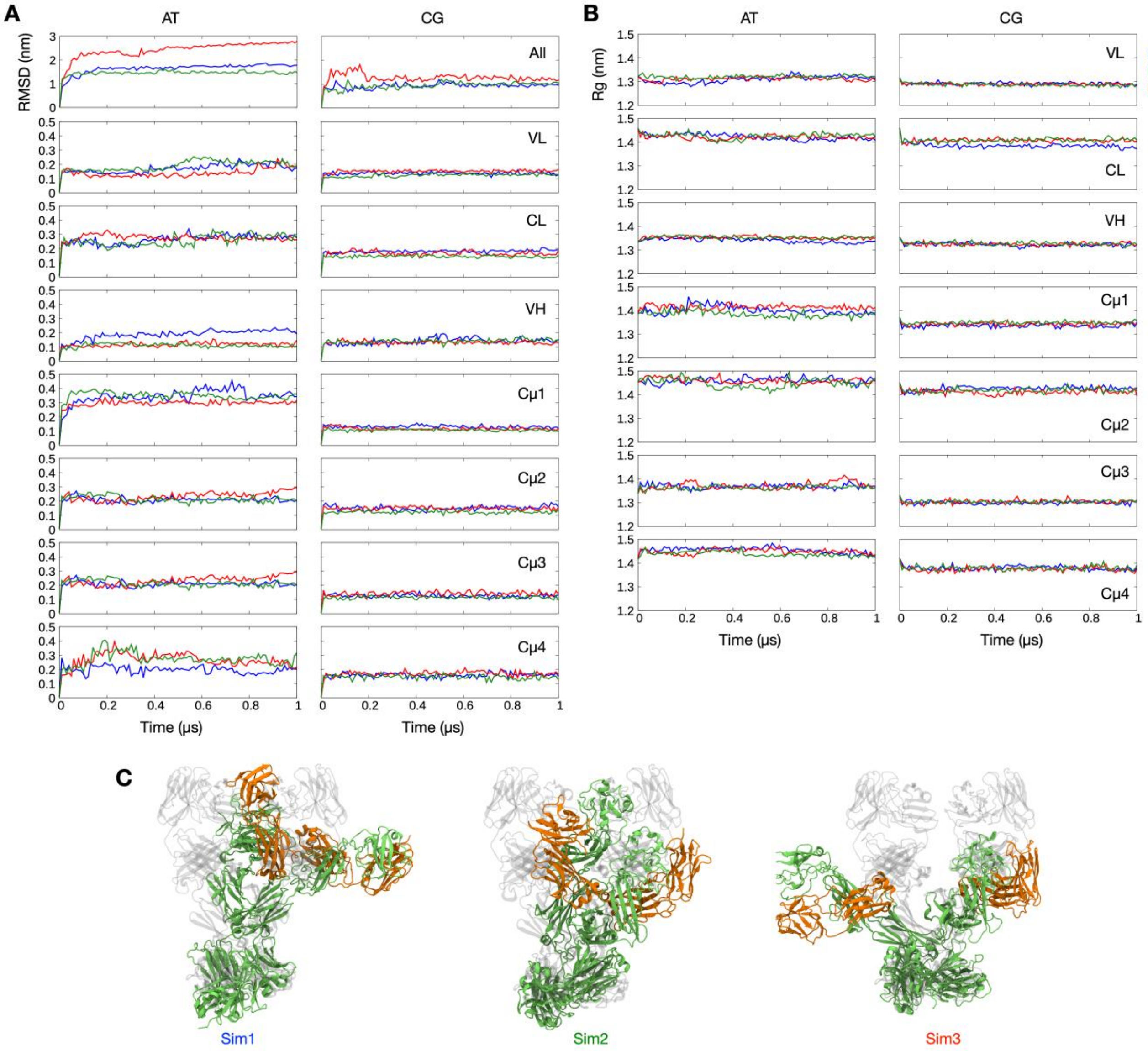


**Figure S2 expositing Figure 1: Structural stability of the homology models of IgM protomers in all-atom and coarse-grained simulations. (A)** Backbone root mean square deviations (RMSD) for Pertuzumab from three independent atomistic (left) and coarse-grained (right) simulations for the whole antibody and each Ig domain. **(B)** The radius of gyration of each Ig domain of Pertuzumab from three independent atomistic (left) and coarse-grained (right) simulations. (C) Snapshots at the end of the three atomistic simulations showing IgM in cartoon representation with heavy chains in green and light chains in orange. The structure of the model at the beginning of the simulation is shown in grey for comparison.

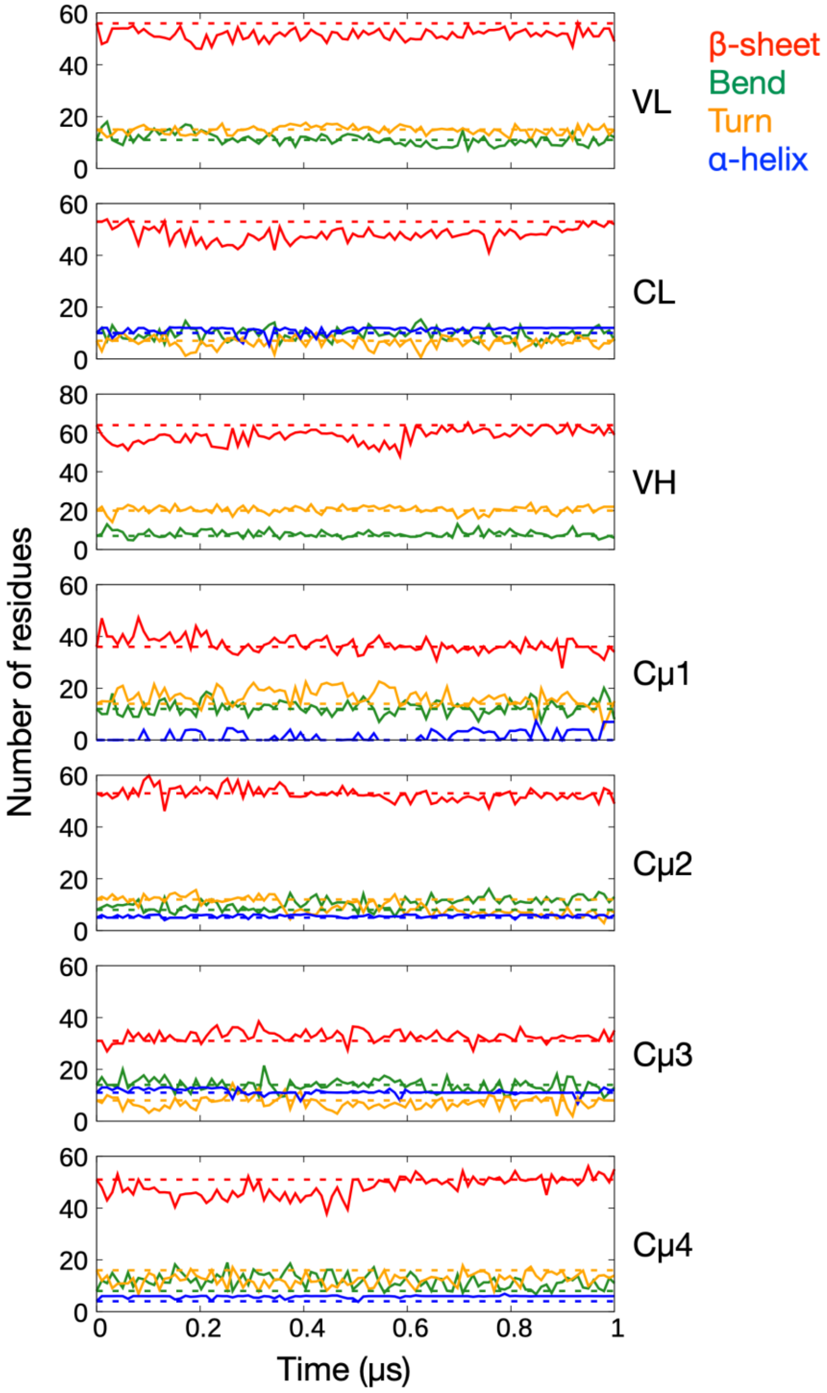


**Figure S3 expositing Figure 1: Secondary structure preservation of each Ig domain in all-atom simulations.** Average number of residues forming β-sheets, bends, turns and α-helices throughout three independent 1 μs simulations. Dotted lines show the number of residues forming these secondary structures at the beginning of the simulations.

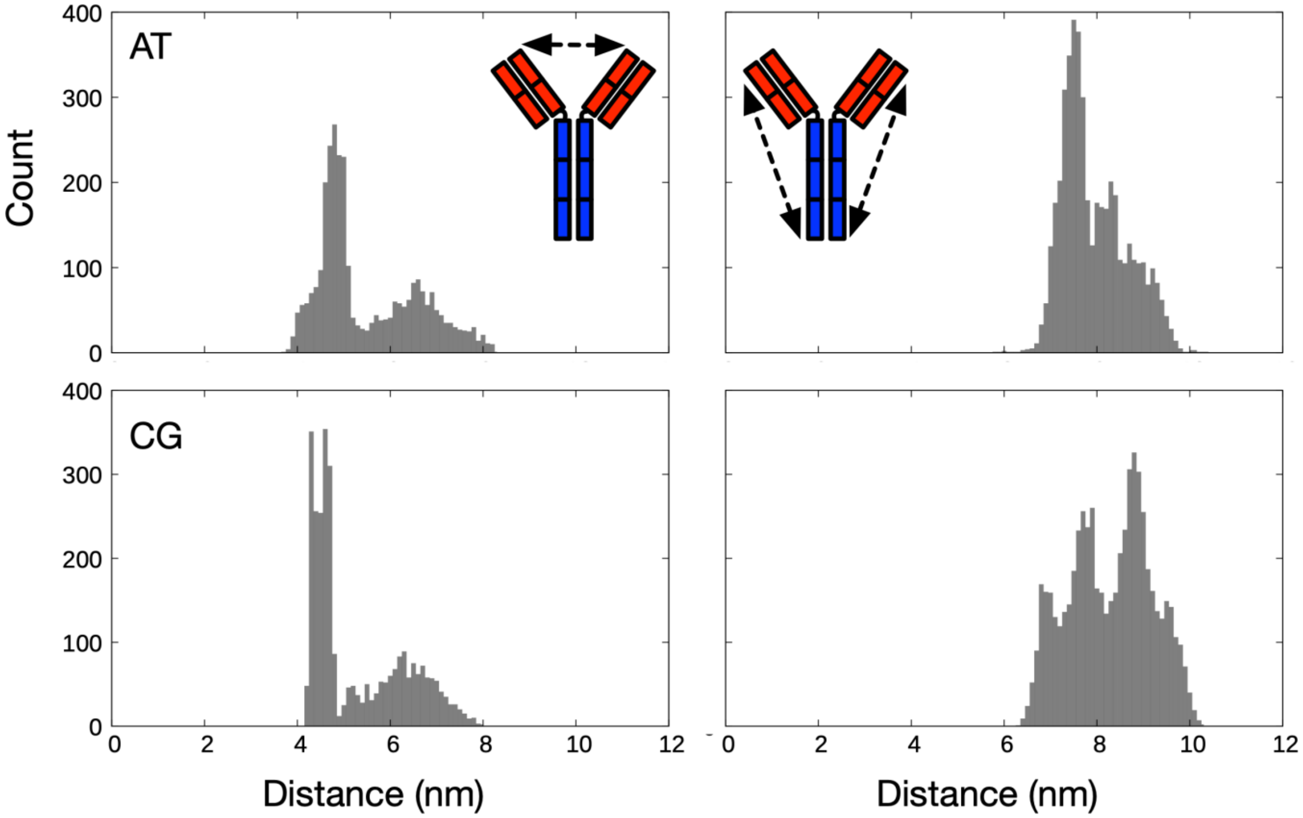


**Figure S4 expositing Figure 1: The distribution of distances from three 1 μs atomistic (top) and coarse-grained (bottom) simulations between the centers of mass of the two Fab domains in Pertuzumab (left), as well as between each Fab domain and the Cμ4 domain (right).**

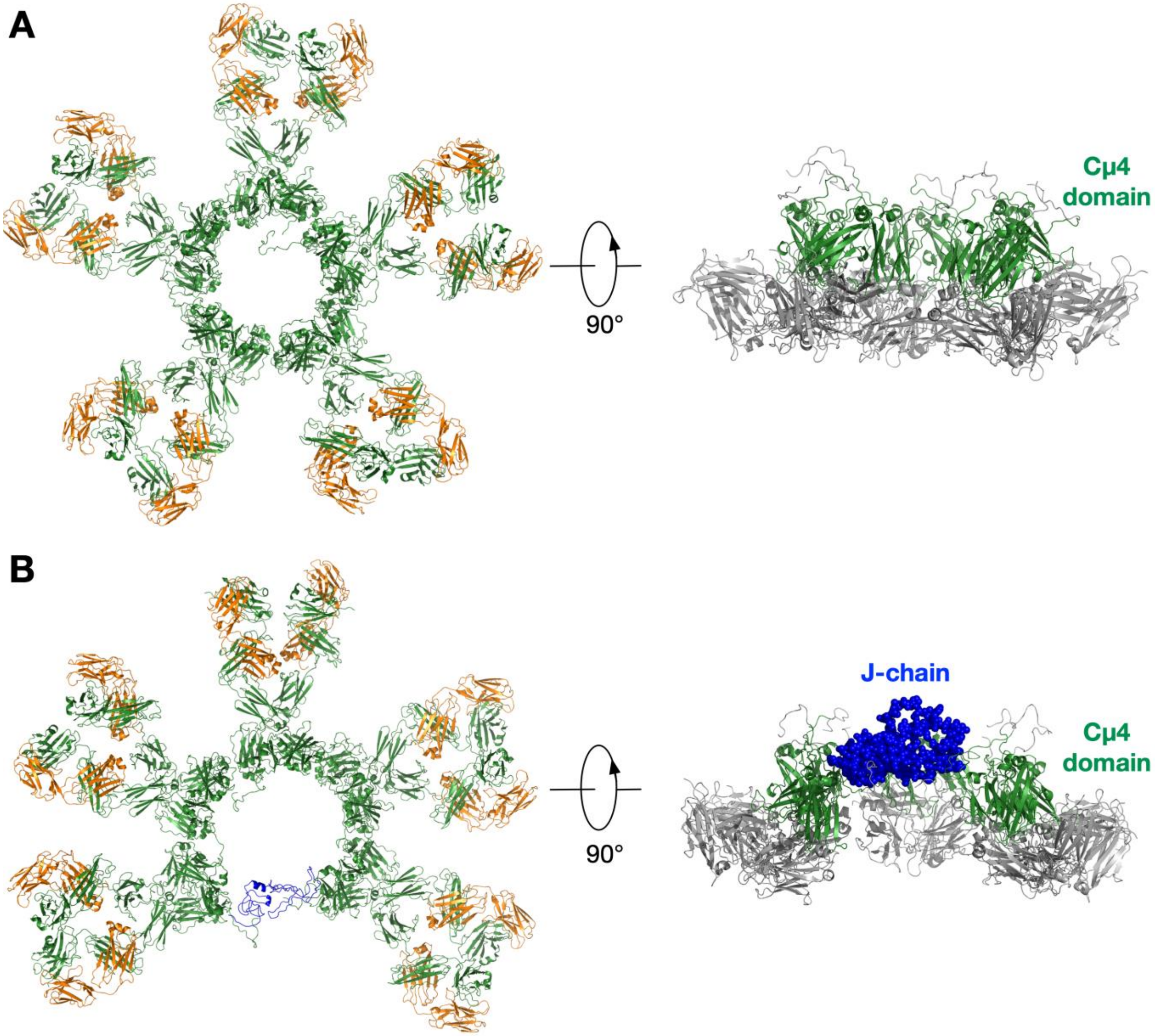


**Figure S5 expositing Figure 1: Pentameric Pertuzumab IgM.** The model is shown **(A)** without and **(B)** with a J-chain in cartoon representation. Heavy chains are in green, light chains are in orange, and the J-chain is in blue. Side views highlighting the protruding Cμ4 domain and the J-chain are shown on the right.

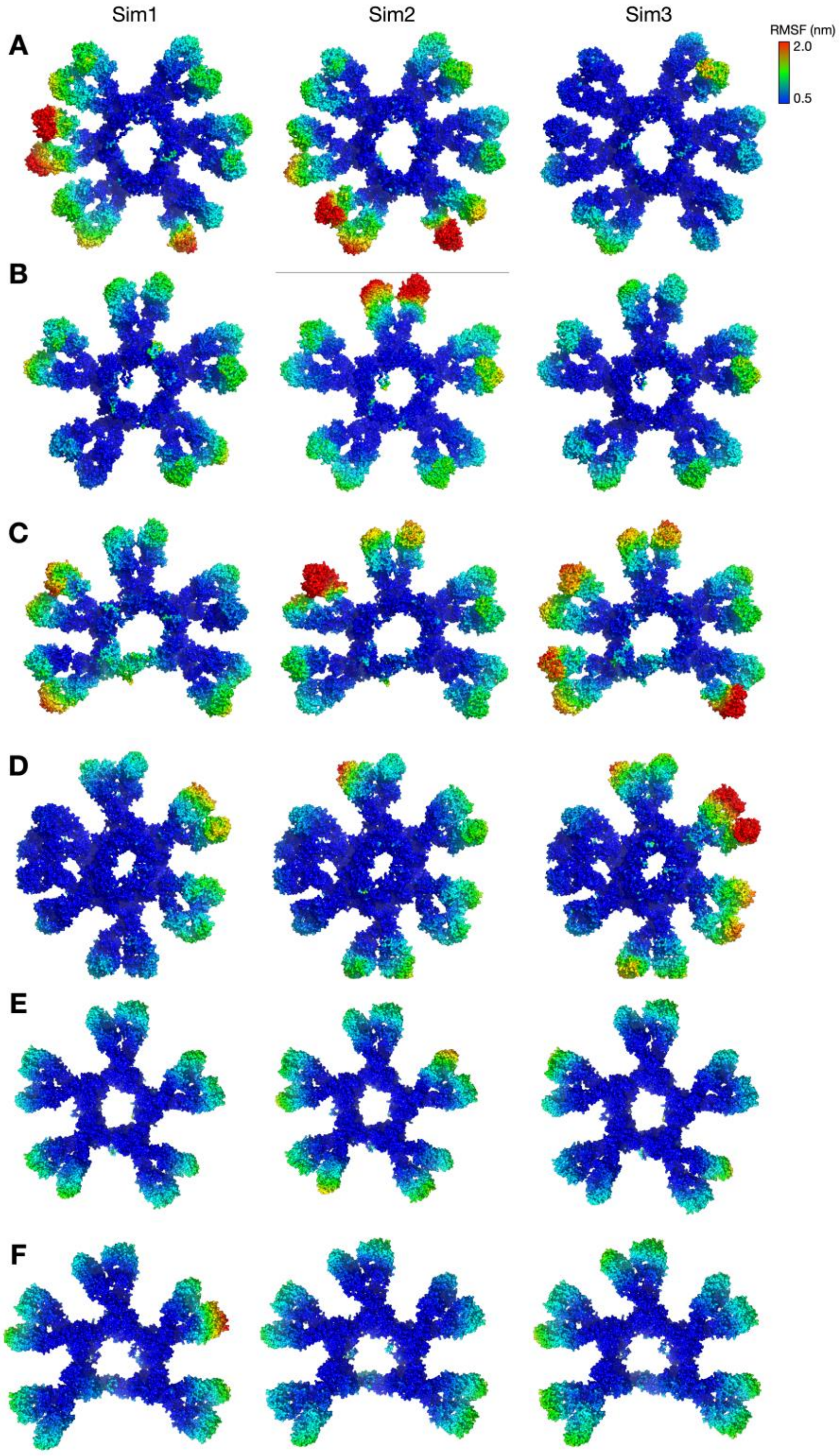


**Figure S6 expositing Figure 2**: **Flexibility of the Fab domain in IgM multimers.** RMSF from three 10 µs simulations values mapped onto the protein surface for: (A) Pertuzumab IgM hexamer; (B) Pertuzumab IgM pentamer; (C) Pertuzumab IgM pentamer with a J-chain; (D) Trastuzumab IgM hexamer; (E) Trastuzumab IgM pentamer; (F) Trastuzumab IgM pentamer with a J-chain.

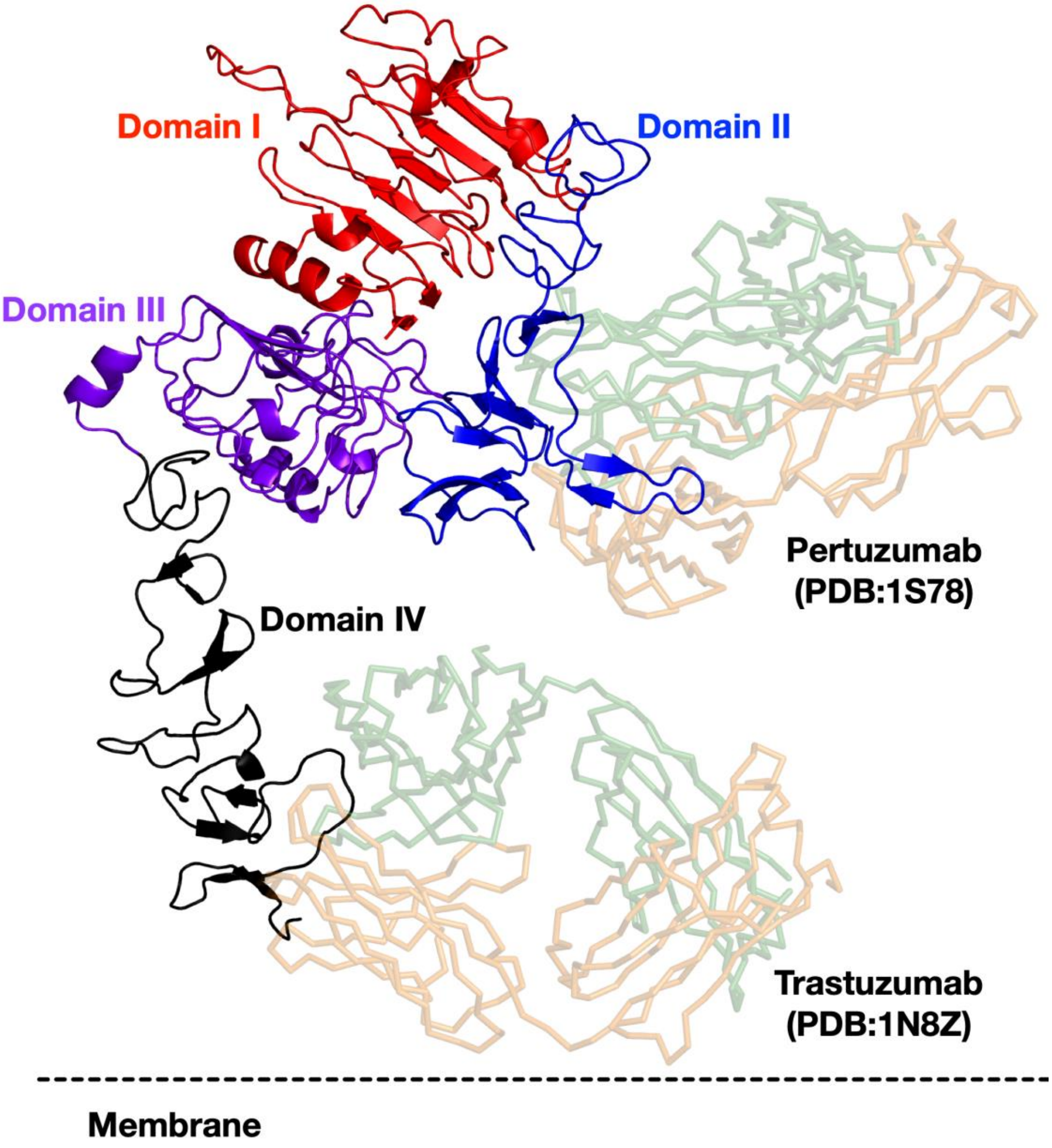


**Figure S7 expositing Figure 4: The binding sites of Pertuzumab and Trastuzumab on HER2 extracellular domain (ECD) as elucidated by X-ray crystallography (PDB: 1S78 and 1N8Z, respectively).** The HER2 ECD is shown in cartoon representation and coloured based on its subdomains. The antibodies are shown in ribbon representation with the heavy and light chains in green and orange, respectively.
